# Selection, recombination and population history effects on runs of homozygosity (ROH) in wild red deer (*Cervus elaphus*)

**DOI:** 10.1101/2023.01.19.524729

**Authors:** Anna M. Hewett, Martin A. Stoffel, Lucy Peters, Susan E. Johnston, Josephine M. Pemberton

## Abstract

The distribution of runs of homozygosity (ROH) may be shaped by a number of interacting processes such as selection, recombination and population history, but little is known about the importance of these mechanisms in shaping ROH in wild populations. We combined an empirical dataset of >3,000 red deer genotyped at >35,000 genome-wide autosomal SNPs and evolutionary simulations to investigate the influence of each of these factors on ROH. We assessed ROH number and location in two populations of red deer (a focal and comparison) to investigate the effect of population history. We investigated the role of recombination using both a physical map and a genetic linkage map to search for ROH. We found differences in ROH distribution between both populations and both map types indicating that population history and local recombination rate have a substantial effect on ROH. Finally, we ran forward genetic simulations with varying population histories, recombination rates and levels of selection, allowing us to further interpret our empirical data. These simulations showed that population history has a greater effect on ROH distribution than either recombination or selection. We further show that selection can cause genomic regions where ROH are common, so called ROH hotspots, only when the effective population size (N_e_) is large or selection is particularly strong. In populations having undergone a population bottleneck, genetic drift can outweigh the effect of selection. We show that most ROH hotspots in the Rum population are in line with expectations from neutral simulations, however two ROH hotspots show possible signatures of selection. Overall, we conclude that in this population, genetic drift resulting from a historical population bottleneck is most likely to have resulted in the observed ROH distribution, with selection possibly playing a minor role.

## Introduction

The inheritance of two genomic segments that are identical by descent due to mating of related individuals generates runs of homozygosity (ROH) in the offspring (Gibson *et al*, 2006). In recent years, with increased availability of high-density genomic data due to SNP arrays and whole genome sequencing, the study of ROH has increased (Ceballos *et al*, 2018a; Curik *et al*, 2014; Pryce *et al*, 2014). Previous studies have shown that the length, abundance and genomic location of ROH vary considerably between populations and can be affected by a number of related processes, including – but not limited to – selection, recombination, and population history (Ceballos *et al*, 2018b; Curik *et al*, 2014; Kardos *et al*, 2017). By investigating ROH, it is possible to gain insights into the relative importance of the mechanisms that cause them.

An allele under strong positive selection is expected to increase in frequency in a population, which will result in increased homozygosity at the site and neighbouring sites and eventually lead to a region where a ROH is common in the population (Sabeti *et al*, 2002). This concept has been widely applied to identify signatures of selection in humans (Nothnagel *et al*, 2010; Pemberton *et al*, 2012), livestock (Ghoreishifar *et al*, 2020; Kim *et al*, 2013; Mastrangelo *et al*, 2017; Peripolli *et al*, 2018; Purfield *et al*, 2017; Zhang *et al*, 2022), and a number of other organisms (Gorssen *et al*, 2021; Grilz-Seger *et al*, 2019; Kumar *et al*, 2020). By searching for regions in the genome where ROH are common, often called ROH hotspots (Pemberton *et al*, 2012) or ROH islands (Nothnagel *et al*, 2010), it is possible to identify genomic regions of interest and even identify genes under selection. For example, in dairy cattle, genes associated with lactation and milk yield are present in regions where more than 50% of the population harbour a ROH (Peripolli *et al*, 2018).

ROH length and position also correlate inversely with local recombination rate (Pemberton *et al*, 2012). Recombination rate varies within and between chromosomes (Johnston *et al*, 2017; Kawakami *et al*, 2014; McVean *et al*, 2004), and studies have shown that ROH hotspots tend to be sites of lower recombination rate (Kardos *et al*, 2017; Stoffel *et al*, 2021). In low recombination regions, long haplotypes are rarely broken up by meiotic crossovers, meaning they are more likely to come together in longer ROH. Conversely, in high recombination regions, long haplotypes are more likely to be broken down, resulting in shorter ROH (Bosse *et al*, 2012). Additionally, the power to detect ROH is higher in regions of low recombination rate, leading to an increased likelihood of identifying ROH, and in turn ROH hotspots (Kardos *et al*, 2017). Recombination rate is not, however, mutually exclusive to selection, and selection may interact with recombination rate to influence ROH distribution (Kardos *et al*, 2017). Selection can reduce genetic variation and effective population size (N_e_) at closely linked loci (via a selective sweep), with the extent of reduced genetic variation dependent on the selection coefficient and local recombination rate (Charlesworth, 2009). It is also important to note the potential for interactions between recombination rate and population history. Species or populations which are closely related are likely to have similar broad-scale recombination rate patterns across the genome, which can again result in more similar distributions of ROHs (Nothnagel *et al*, 2010).

Intuitively, population history events such as a population bottleneck or effective migration, can directly influence ROH abundance (Ceballos *et al*, 2018b; Foote *et al*, 2021; Mooney *et al*, 2021). The more related members of a population are, the more inbreeding will occur and the more ROH will be present. Individuals from the same populations tend to have a similar inbreeding coefficients, number of ROHs and distribution of ROH lengths, reflecting their shared population history (Bertolini *et al*, 2018; Bosse *et al*, 2012; Cardoso *et al*, 2018). A more specific consequence of population history can occur following a population bottleneck, where drift and selection play an important role in shaping the ROH landscape. Genetic drift may result in certain alleles becoming fixed by chance, increasing the likelihood of generating ROHs at a site under inbreeding. Moreover, the interaction between effective population size (N_e_) and selection may be key in the formation of ROH hotspots. The product of N_e_ and the selection coefficient (*s*) of a new mutation can determine its likelihood of fixation in the population (Falconer, 1983). When *s* is high, an allele is likely to become fixed; however, when N_e_ is small, genetic drift can outweigh the force of selection (Petit and Barbadilla, 2009).

Together, selection, recombination rate and population history are likely to play interacting roles in shaping the incidence and genomic locations of ROH in a population. However, few studies have investigated all these factors in a wild population. Here, we aimed to investigate the influence of each factor on ROH number and location in a wild population of red deer inhabiting the island of Rum, Scotland. This individually-monitored population has experienced a population bottleneck, shows evidence for inbreeding depression (Huisman *et al*, 2016) and has a detailed genetic linkage map (Johnston *et al*, 2017) making it an ideal population to address our aims.

## Methods

### Study populations

The red deer population inhabiting the north block of the Isle of Rum, Scotland (57°0’N, 6°20’W) has been studied at an individual level since 1971 (Clutton-Brock *et al*, 1982), and is the main focus of this study. DNA was extracted from ear punches from calves captured soon after birth or darted adults, post-mortem tissue or cast antlers (See Huisman *et al* (2016) for full details). DNA samples were genotyped at 50,541 attempted SNP loci on the Cervine Illumina 50K BeadChip (Brauning, 2015). SNP genotypes were clustered and scored using Illumina GenomeStudio v2.0, and were subject to quality control with the following parameters: SNP minor allele frequency (MAF) > 0.001, ID genotyping success > 0.9, and SNP genotyping success > 0.99. In addition, SNPs mapped to the sex chromosomes were removed. This resulted in a final data set of 39,587 autosomal SNPs genotyped in 3046 individuals (Johnston *et al* (2017).

This study also used equivalent genotype data for 157 individuals from a mainland population of red deer from Argyll, Scotland. Details of sample collection and DNA extraction can be found in McFarlane *et al* (2020). Samples were genotyped as above and quality control was carried out by McFarlane *et al* (2020) as above with the exception of SNP genotyping success >0.9. These individuals were originally genotyped as part of a study of red x sika deer (*Cervus nippon*) hybridisation and were pure red deer according to that study (McFarlane *et al*, 2020).

### Calling runs of homozygosity

We searched for runs of homozygosity in each genotyped individual using the --*homozyg* function in PLINK v1.90 (Purcell *et al*, 2007). We used two different estimates of marker positions to call ROH: physical distances in base pairs (bp) and genetic distance in centimorgans (cM): the genetic map accounts for variable recombination rate through the genome, whereas the physical map is independent of recombination (Kardos *et al*, 2017). Recombination distance was based on a genetic linkage map previously constructed for the Rum deer population using 38,038 SNP markers and pedigree information (Johnston *et al*, 2017). Physical marker positions were obtained from the red deer (*Cervus elaphus*) genome assembly version mCerEla1.1 which was constructed using DNA from a female red deer originating from Rum (Pemberton *et al*, 2021). For consistency between searches, we assumed 1Mb ≈ 1cM (Actual value: 1Mb = 1.04cM; (Johnston *et al*, 2017)). The following parameters were used to identify ROH in both cases: --*autosome-num* 33 (a flag to specify the 33 autosomes in red deer), --*homozyg-snp* 40 (minimum number of SNPs in a ROH), --*homozyg-kb* 2500 (minimum length of a ROH in kb), --*homozyg-density* 70 (minimum density, 1 SNP per 70kb), --*homozyg-window-snp* 35 (sliding window size), --*homozyg-window-missing* 4 (number of missing SNPs allowed in a window), *homozyg-het* 0 (number of heterozygote SNPs allowed in a ROH), and --*maf* 0.01 (minor allele frequency threshold). Following the use of these parameters in PLINK 35,132 autosomal SNPs and 3046 individuals were analysed in the Rum dataset and 34,673 autosomal SNPs and 157 individuals in the Argyll dataset.

We used simulations (details given below) to show that the chosen search parameters for our average SNP density can capture the ‘true ROH’ present (Supplementary Figure 1).

### Estimate of inbreeding coefficient

We used the called ROH to estimate individual genome-wide inbreeding coefficient, F_ROH_ (McQuillan *et al*, 2008), using the sum of Mb or cM in ROH across all autosomes divided by the total Mb of autosomes estimated as 2591.86Mb from the genome assembly mCerEla1.1. As above we assumed 1Mb ≈ 1cM (Johnston *et al*, 2017), therefore the same value was used as the denominator in both instances.

### ROH Hotspots

Following our ROH search, we determined ROH hotspots across the genome by estimating the proportion of individuals with ROH covering a SNP locus i.e. the ROH density. ROH density was calculated as a SNP-by-SNP measure as follows:

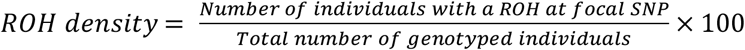

We used the 99^th^ percentile value for ROH density as the hotspot threshold, such that any SNP with a value equal to or above this threshold was classed as a hotspot SNP. This outlier/percentile approach has been used by a number of recent publications (Bertolini *et al*, 2018; Cesarani *et al*, 2022; Mastrangelo *et al*, 2018; Purfield *et al*, 2017; Zhang *et al*, 2022).

We found that ROH density was positively correlated with the density of SNPs such that fewer ROH were identified in regions where fewer SNPs were genotyped. This issue with the PLINK algorithm for calling ROH has previously been identified for this type of analysis (Nandolo *et al*, 2018). Windows of size 1500Kb (or equivalent in cM) and sliding 100Kb (or equivalent) allowed for the assessment of the relationship between number of SNPs in a window and number of ROH found. Following assessment of both populations using the bp and cM datasets, we chose a minimum density of 23 SNPs per 1500kb to minimise the correlation, but maximise consistency between methods and the number of SNPs remaining in the analysis (Supplementary Figure 2, Supplementary Table 1). All SNPs within windows that fell below this density were discarded from the analysis of hotspots. Additional post-ROH-search quality control removed the first and last 40 SNPs from each chromosome to account for the fact that fewer ROH will have been called in these regions as a ROH cannot span past the ends of a chromosome. (See Supplementary Table 1 for number of remaining SNPs post-ROH search)

### Haplotype diversity

As an additional analysis to detect possible signatures of selection through a drop in genetic diversity, we estimated haplotype diversity for the Rum dataset and in our simulated data. Phased autosomal haplotypes for each individual in the Rum dataset were obtained using AlphaPeel v0.0.1 (Whalen *et al*, 2018). Phasing was conducted using the red deer pedigree and genomic dataset, using multi-locus peeling, which takes into account that nearby loci are more likely to be inherited jointly from a parental haplotype. Five peeling cycles were run, and only phased genotypes with a probability of > 0.95 were retained. Phasing success was very high overall, with an individual mean phasing rate (i.e., the proportion of an individual’s genotypes that were phased) of ~99% in individuals born after 1980.

Phased haplotypes were then split into 20 SNP windows with a 10 SNP sliding increment and any window with missing calls within it was removed. The number of unique 20-SNP haplotypes in each window and the number of occurrences of each, was counted. A haplotype diversity measure was based on Simpson’s diversity index for measure of biodiversity:

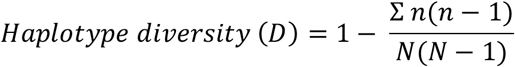

Where *n* = the number of occurrences of each haplotype and *N* = total number of occurrences of all haplotypes and the range of *D* is from 0 (low diversity) to 1 (high diversity).

### Simulations

Simulations were carried out in SLiM v3 (Haller and Messer, 2019) to test the effect of selection, recombination rate and population history on the distribution of ROH across a simulated 100Mb chromosome. Each simulation model was run for 23 iterations. We used three different population history scenarios: one reflecting the Rum population, one with no population bottleneck and one with a more severe population bottleneck than the Rum population (Figure 1). All scenarios started with an effective population size of 7500 set to run for 75,000 generations as a burn-in (10*Ne) (Haller and Messer, 2019). This starting effective population size was based on N_e_ = ~1/10 census population size (Frankham, 2007). An estimation of the red deer population in Scotland at the beginning of the 20^th^ century was ~150,000 individuals, which has since doubled to ~300,000 at the start of the 21^st^ century (Pepper *et al*, 2020). Assuming a similar rate of population increase, the red deer population of Scotland (and the rest of the UK) was ~75,000 at the beginning of the 19^th^ century.

**Figure 1.**
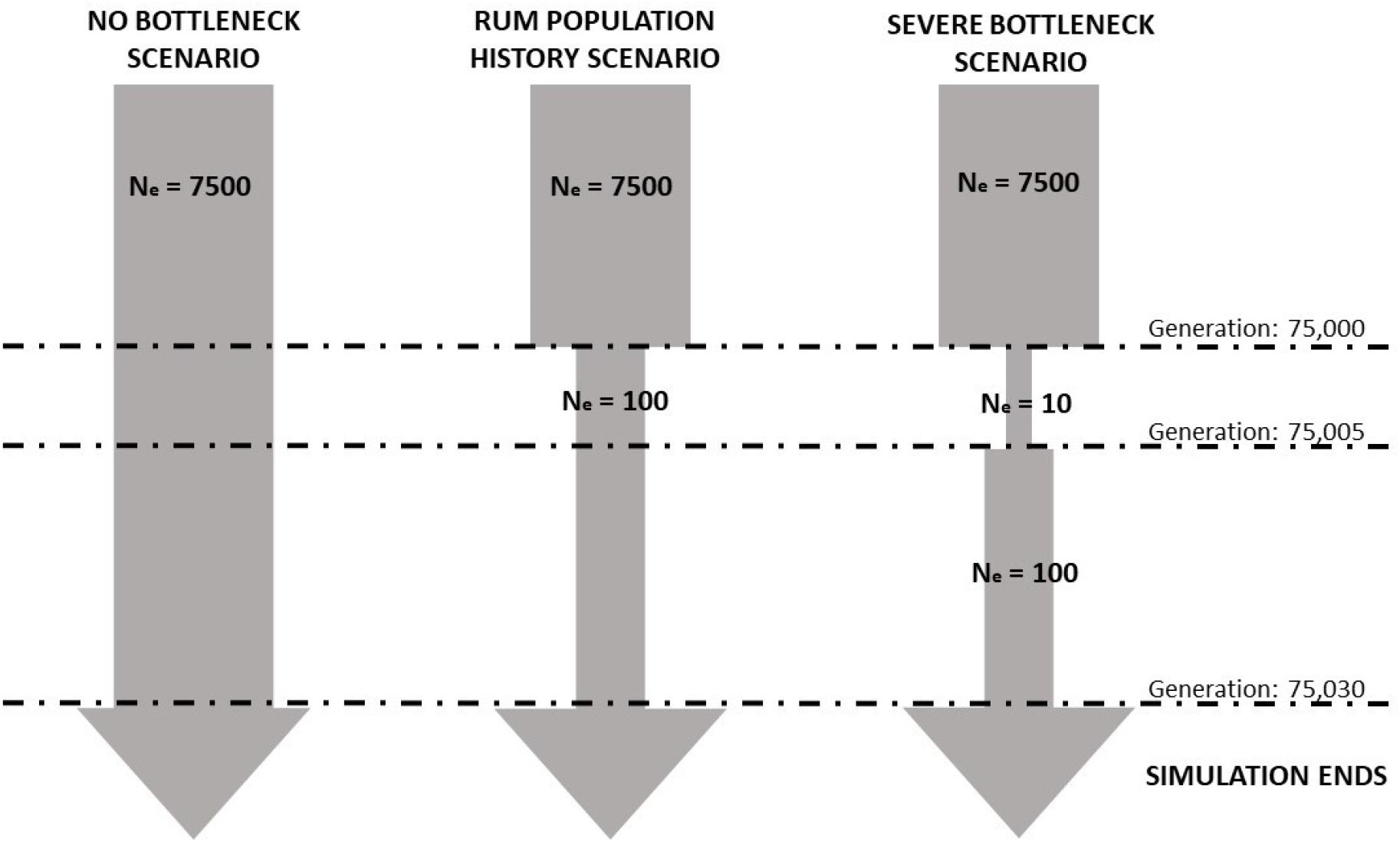
Representation of model design for three simulated population histories. All simulations begin with an effective population size (N_e_) of 7,500. Dashed lines show the time points when changes occur to the effective population size, with the generation shown on the right. All simulations ended at generation 75,030.

The population history of Rum is somewhat unknown, as an unrecorded number of individuals were introduced from various red deer herds in Scotland and England beginning in 1845 (Marshall, 1998). Therefore, to simulate this bottleneck event in the Rum population history we dropped N_e_ to 100 at generation 75,000. The simulation ended at generation 75,030 (simulated present day) with a current N_e_ of 100 individuals reflecting the current population of around 1000 individuals. In the more severe bottleneck scenario we dropped N_e_ to 10 at generation 75,000, and then increased it to 100 at generation 75,005. As above the simulation ended at generation 75,030 with 100 individuals. The no bottleneck scenario maintained an N_e_ of 7,500 until generation 70,030 (Figure 1). The output for all scenarios was the genomic data for the last simulated generation.

Four different models were tested for each population history scenario: 1) neutral; 2) varied recombination rate; 3) varied recombination rate plus selection; and 4) varied recombination rate plus stronger selection. All models had a mutation rate of 1×10^-8^ per base pair per generation (Kyriazis *et al*, 2021).

Model 1 was a neutral model set to have a constant recombination rate across the chromosome at 1.038 cM/Mb based on estimates from the linkage map (Johnston *et al*, 2017), with every mutation set as neutral (i.e. selection coefficient = 0).

Model 2 had a varied recombination rate across the chromosome. All deer chromosomes, with the exception of chromosome 5, are acrocentric (i.e. the centromere occurs very close to the end of the chromosome) and Johnston et al (2017) demonstrated that recombination rate shows consistent broad scale variation across chromosomes in red deer. For example, sex-averaged recombination rates are higher in peri-centromeric regions (0 – 25% of the chromosome length) in comparison to the rest of the chromosome where recombination rates are more constant, with a slight elevation towards the sub-telomere. We split the simulated 100Mb chromosome into 10 Mb regions and applied a recombination rate to each region according to this observed data. For regions 1-10 the recombination rates were as follows: 1.75 cM/Mb, 1.23 cM/Mb, 0.89 cM/Mb, 0.81 cM/Mb, 0.74 cM/Mb, 0.80 cM/Mb, 0.87 cM/Mb, 1.05 cM/Mb, 1.20 cM/Mb, and 0.67 cM/Mb.

Model 3 incorporated neutral mutations (fixed selection coefficient = 0), beneficial mutations (mean selection coefficient = 0.001 drawn from gamma distribution with shape parameter = 0.2 and a dominance coefficient = 0.5 i.e. additive), and deleterious mutations (mean selection coefficient = −0.01 drawn from gamma distribution shape parameter 0.2, and dominance coefficient = 0.1 under the assumption deleterious mutations are partially recessive). Neutral, beneficial and deleterious mutations occurred in the ratio 3:1:10 respectively (Huber *et al*, 2017; Kim *et al*, 2017; Kyriazis *et al*, 2021). This model also included varied recombination as specified for Model 2.

Model 4 included variable recombination as in model 2 but higher selection coefficients than in model 3. We increased the selection coefficients by a factor of five, i.e. beneficial mutations had a mean selection coefficient = 0.005 and deleterious mutations had a mean selection coefficient = −0.05).

Following simulation runs 100 (and 7,500 for the no bottleneck scenario) individual genomes were output from SLiM and ROH were called with parameters as described above without the minor allele frequency threshold. In addition, we carried out haplotype diversity calculations on these outputs as described above, further details given in Supplementary Figures 7 & 8. All SLiM scripts are available at https://github.com/annamayh/ROH_hotspots_manu.

## Results

### ROH abundance and length between populations

Across all 3,046 Rum deer, we found 68,643 ROH using the physical map (bp positions) and 45,814 ROH using genetic map (cM positions). In the 157 Argyll deer, the equivalent figures were 1,825 and 1,227 ROH, respectively. The Rum population had significantly more ROH per individual and higher inbreeding coefficients (F_ROH_) than the Argyll population (Table 1). The majority of ROH we found were between 2.5 Mbp (minimum ROH length included) and 16 Mbp, with a mean ROH length of ~6 Mbp; a small number of individuals had ROH > 16Mbp (Table 1, Supplementary Table 2).

**Table 1.** Comparison of mean ROH length +/- standard deviation (SD), Mean (+/- SD), minimum and maximum number of ROH per individual, mean inbreeding coefficient (+/-SD) and maximum individual inbreeding coefficient in two red deer populations and using two maps to search for ROH.

| | Mean ROH<br>length<br>(Mbp/McM)<sup>1</sup> | Mean<br>number<br>of ROH<br>per ID<sup>2</sup> | Min<br>number<br>of ROH | Max<br>number<br>of ROH | Mean<br>$F_{ROH}$<sup>3</sup> | Max<br>$F_{ROH}$ |
| --- | --- | --- | --- | --- | --- | --- |
| **Rum**<br>**(Physical)** | 6.45 ± 5.4 | 22.54 ±<br>6.0 | 0 | 45 | 0.056<br>±0.02 | 0.300 |
| **Rum**<br>**(Genetic)** | 6.10 ± 4.7 | 15.04 ±<br>4.5 | 0 | 34 | 0.035 ±<br>0.02 | 0.190 |
| **Argyll**<br>**(Physical)** | 6.47 ± 5.9 | 11.62 ±<br>5.3 | 1 | 31 | 0.029 ±<br>0.02 | 0.143 |
| **Argyll**<br>**(Genetic)** | 6.21 ± 5.2 | 7.82 ±<br>4.2 | 0 | 22 | 0.019 ±<br>0.01 | 0.103 |

<sup>1</sup> Difference between map positions used within each population were only significant in the Rum dataset. One-way ANOVA of map on ROH length, p-values: Rum dataset <2.2e-16 and Argyll dataset <0.1
<sup>2</sup> Difference between map positions used within each population were significant. One-way ANOVA of map on number of ROH per ID, p-values: Rum dataset <2.2e-16 and Argyll dataset <9e-12. Differences between populations within maps were significant. One-way ANOVA of population on number of ROH per ID, p-value <2.2e-16 for both maps.
<sup>3</sup> Difference between map positions used within each population were significant. One-way ANOVA of map on $F_{ROH}$ , p-values: Rum dataset <2.2e-16 and Argyll dataset <3e-7. Differences between populations within maps were significant. One-way ANOVA of population on $F_{ROH}$ , p-value <2.2e-16 for both maps.

### ROH hotspots

As summarised above, we found that the marker positions used (bp or cM positions) significantly affected the number of ROH found in the populations (Table 1). We next explored this map effect on the genomic distribution of ROH in the Rum population. We found eight regions on six different chromosomes that were classed as ROH hotspots using the physical map positions, whereas only five regions were ROH hotspots using the genetic map positions, all of which were previously identified using the physical map positions (Figure 2, Table 2, ROH hotspot positions can be found in Supplementary Table 4). No identified hotspot SNPs were shared between the Rum and Argyll populations. (Supplementary Figure 3, Table 2, Supplementary Tables 3 & 4).

**Figure 2.**
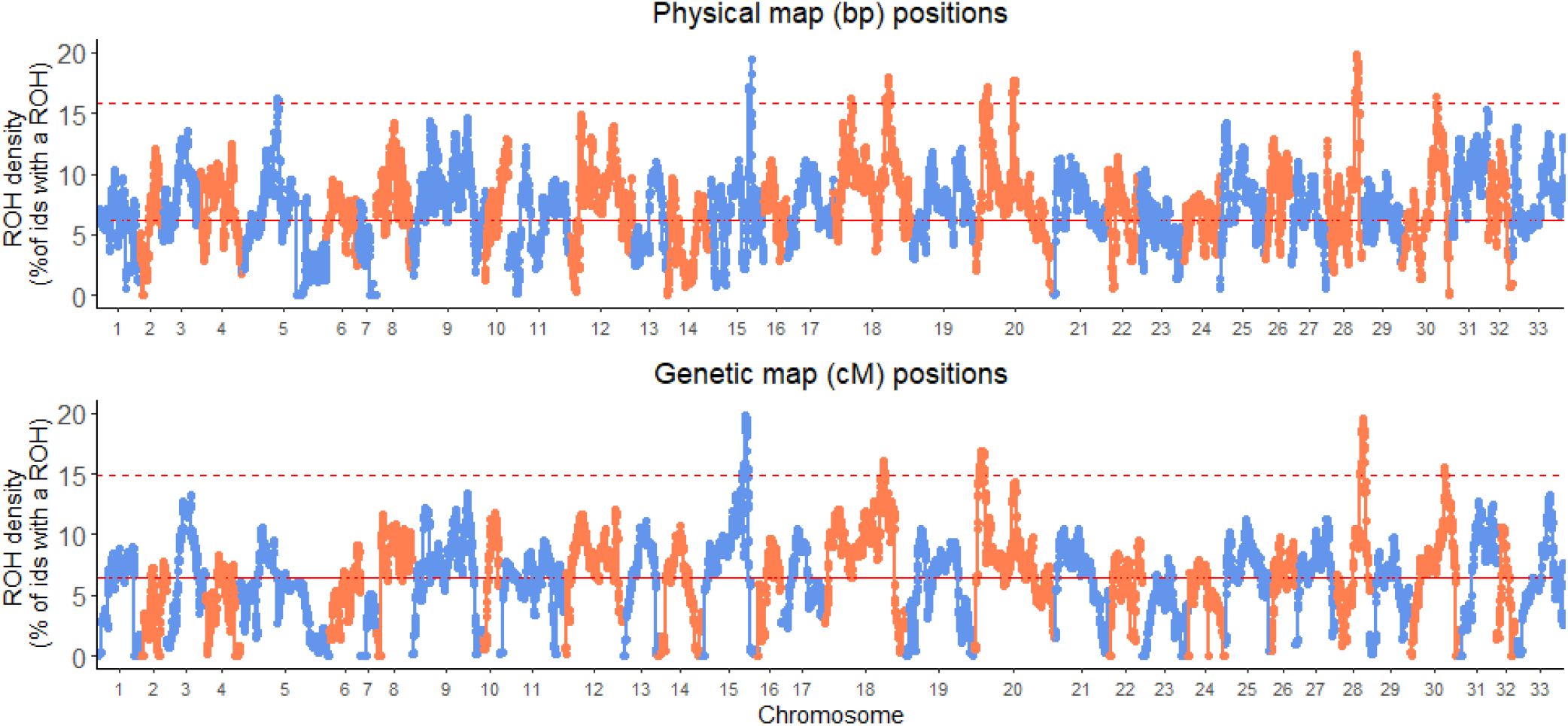
Percentage of Rum individuals with a ROH at each SNP (ROH density) across all chromosomes. The top panel shows the distribution of ROH density using the physical map positions (bp) and the bottom panel shows the same when using genetic map positions (cM). The upper dashed lines indicate the ROH hotspot threshold (99^th^ percentile ROH density) and the solid red line indicates the mean ROH density.

**Table 2.** Comparison of ROH hotspots using two maps to search for ROH. The table shows the mean ROH density, with standard deviation (SD)and the ROH hotspot threshold. Any SNP with a value of ROH density above this threshold value is classed as a ROH hotspot SNP. The table also shows the number of SNPs classed as a ROH hotspot SNP and the chromosomes containing ROH hotspots. The notations (a), (b) indicate independent hotspots on the same chromosomes.

| Map Position used | Mean ROH density ± SD (%) | ROH Hotspot threshold (99<sup>th</sup> percentile ROH density) | # of SNPs over ROH hotspot threshold | Chromosomes containing ROH hotspots |
| --- | --- | --- | --- | --- |
| **Physical** | 6.2 ± 0.02 | 15.9% | 289 | 5,15,18(a),18(b),20(a),20(b),28,30 |
| **Genetic** | 6.4 ± 0.02 | 14.9% | 257 | 15,18(b),20(a),28,30 |

We also looked for ROH hotspots using only short ROH, between 2.5Mb-5Mb. As above, we found a differences between the maps used. We also find the same three ROH hotspots on chromosomes 15, 18 and 28 when using the genetic map positions, plus four other loci reaching just over the threshold (Supplementary Figure 4).

### Haplotype diversity

A low haplotype diversity indicates that certain haplotypes are more common than others, whereas high haplotype diversity indicates haplotypes occur in equal number. We found a sharp decrease in haplotype diversity in two of the five ROH hotspots, on chromosomes 15 and 28, Figure 3. This sharp decrease is consistent for chromosome 15 when different window sizes are used (supplementary Figures 5 & 6). However the diversity decrease on chromosome 28 appears more dependent on window size, with a larger window size showing a less dramatic decline (supplementary Figures 5 & 6). In addition, we show that our neutral simulations for the Rum scenario show minimal drops in haplotype diversity (Supplementary Figure 7). On the other hand, simulations of the Rum scenario including strong selection have multiple drops in haplotype diversity (Supplementary Figure 8), presumably due to the imposed selection coefficients. Moreover, some, but not all, of these drops coincide with ROH hotspots.

**Figure 3.**
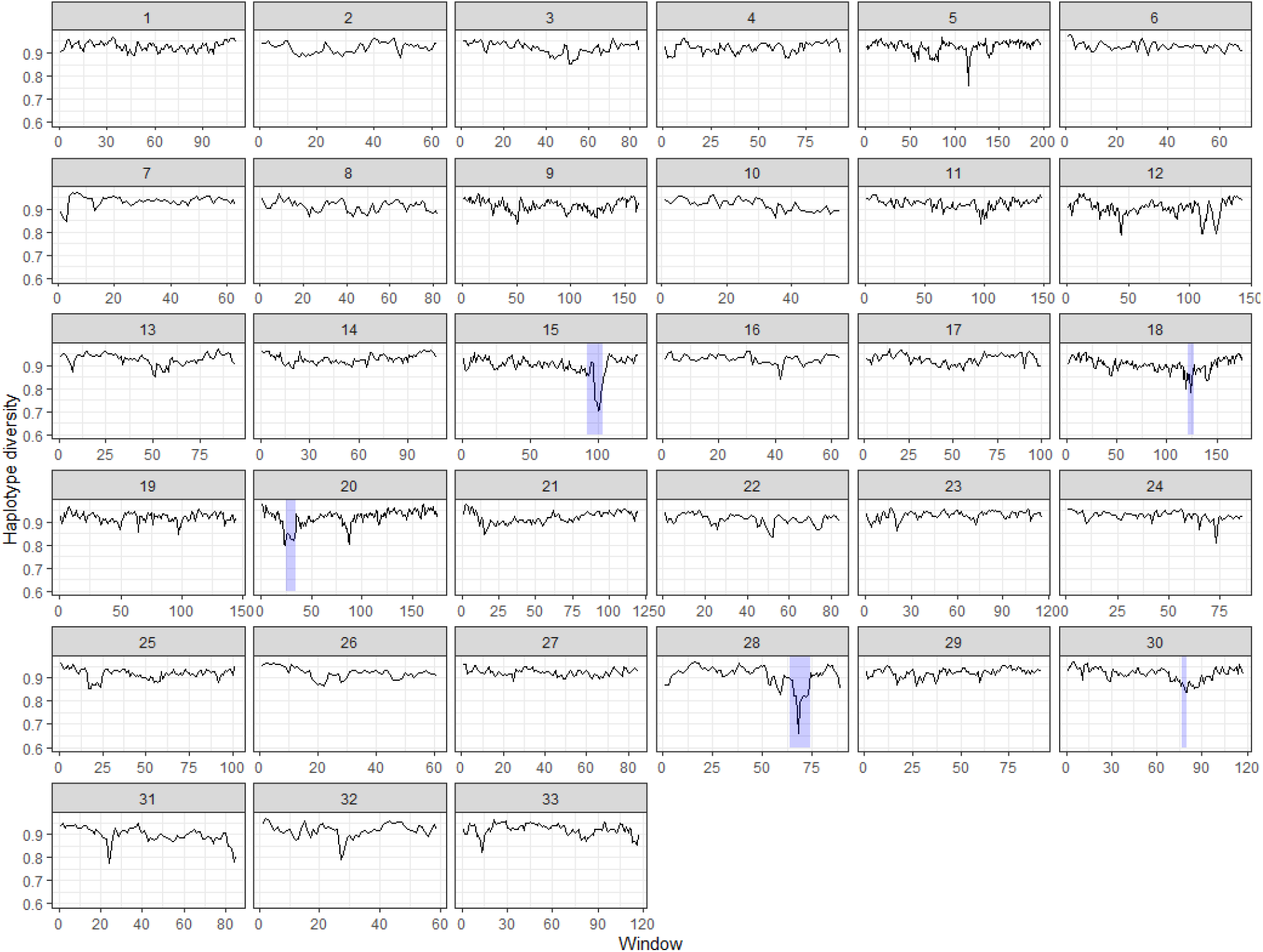
Haplotype diversity for 20-SNP windows in 10 SNP sliding increments across 33 autosomes in the Rum population of red deer. Low haplotype diversity indicates that certain haplotypes are more common; high haplotype diversity indicates haplotypes occur at more even frequencies. ROH hotspot regions are shaded in purple.

### Simulations

In our simulations, we ran 23 iterations of four model designs within three different population histories. In Figure 4, we show the ROH hotspot threshold (Figure 4A) and maximum ROH density (Figure 4B) for each of the 23 iterations, grouped by model design. For each population history scenario, we compared various models to a neutral model with a constant recombination rate throughout the chromosome and no selection. Comparing to this neutral model (Model 1), we investigated the effects of varying recombination rates over the chromosome alone (Model 2), coupled with weak selection (Model 3), or coupled with stronger selection (Model 4).

**Figure 4.**
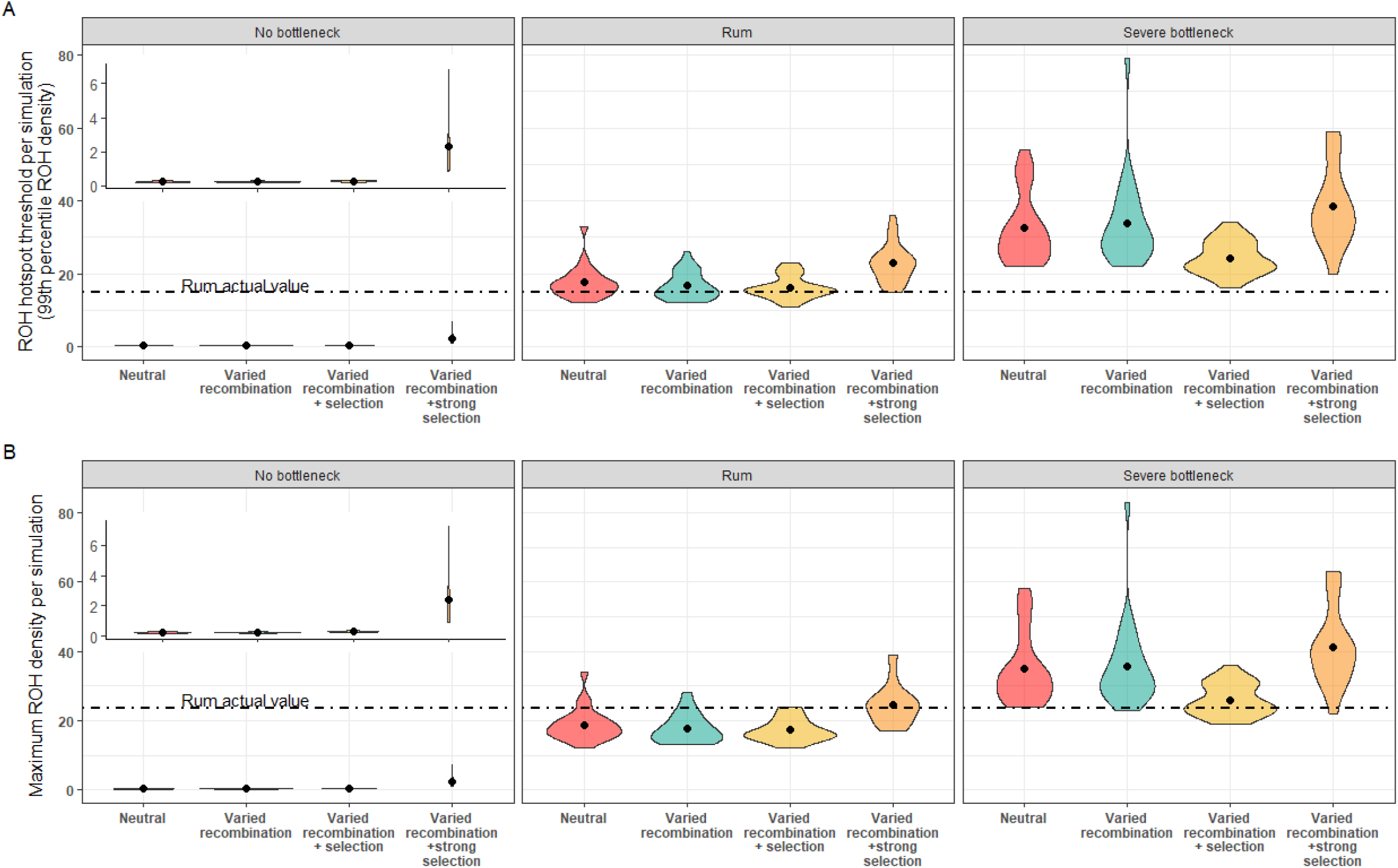
Violin plots showing A) ROH hotspot threshold and B) Maximum ROH density for every iteration. Black dots indicate the mean value for 23 iterations of a simulation, the width of the violin indicates the number of simulations at that value. Each box contains a different simulated population history, from left to right: no historical population bottleneck, Rum population history, severe historical population bottleneck. Each population history was modelled under four conditions: a neutral model with a constant recombination rate and no selection, a model including varied recombination rate, a model including varied recombination and selection and a model including varied recombination rate and strong selection (see Methods for details of parameter values). Inset plot shows a zoomed view of the no-bottleneck scenario. Dashed line show the equivalent value for the empirical dataset.

We found that of all the parameters tested, population history had the greatest effect on ROH. The mean number of ROH per individual was 0.06, 1.72 and 3.25 for the no bottleneck, Rum and severe bottleneck scenarios, respectively (note that these values are for a single simulated 100Mb chromosome), and the average Kb in a ROH per individual was 260Kb, 9,091Kb and 15,158Kb, respectively (Supplementary Table 5). Moreover, when we compared ROH density between population histories (compare across panels in Figure 4) there was noticeable variation. As expected, these differences in ROH patterns between different scenarios reflected the expected level of inbreeding and N_e_ in the simulated populations. Additionally, the severe bottleneck model has the most variation between iterations, further highlighting the impact population history, particularly a bottleneck, can have on ROH.

We found little impact of varied recombination rate or weak selection on ROH. In all population histories, Models 2 and 3 did not differ greatly from the neutral Model 1 (Figure 4). On the other hand, strong selection did have a substantial effect, particularly in the no bottleneck scenario. Model 4 showed higher maximum ROH density and hotspot threshold values (Figure 4). This pattern is most visually obvious in the no bottleneck scenario and least in the severe bottleneck scenario (Figure 4 left and right panels respectively).

In order to assess the likelihood that ROH hotspots identified in the Rum population may be caused by selection, we compared the observed data to our simulated data. We show that results from Model 3, including weak selection, are not appreciably different from the neutral model and both overlap with the empirical values (Figure 4A middle panel, Supplementary Figure 9). Model 4, including strong selection, does not overlap with the empirical data for the ROH threshold, with the empirical value for this falling outside the 95% confidence intervals (Figure 4A middle panel, Supplementary Figure 11). This indicates that most ROH hotspots in the Rum dataset falling over this threshold are not caused by strong selection. However, the maximum ROH density in the Rum dataset is within the 95% confidence intervals from Model 4 and coincides with the mean across iterations (Figure 4B middle panel, Supplementary Figure 10), suggesting that the maximum ROH hotspot peak(s) seen in Figure 2 could be areas of strong selection. However, this maximum value also overlaps with the confidence intervals of other models (Supplementary Figure 10).

## Discussion

### Population history and inbreeding level

In this study, we investigated the roles of population history, recombination and selection on the number and distribution of ROH in a wild red deer population, and combined this with forward genetic simulations. We first found that the Rum population had a higher level of inbreeding than the comparison Argyll population, and this difference was consistent between methods (Table 1). This difference probably reflects the different recent histories of the two populations. The Rum population became isolated from the mainland ~150 years ago, has had few introductions since and currently numbers around 1,000 individuals, therefore is more likely to inbreed, while the Argyll population is continuous with the rest of the Scottish mainland population which numbers in the hundreds of thousands. The direct influence of population history on ROH number has been previously documented in a number of wild species (Foote *et al*, 2021; Nguyen *et al*, 2022), including in a study of red deer which included Rum deer samples (de Jong et al. 2020).

Of all the variables tested within our simulations, population history had the greatest influence on ROH distribution, reflecting the level of inbreeding in the simulated populations (Figure 4). For example, the simulations with a severe bottleneck had the highest number of ROH as a result of a smaller Ne, reflecting observed results from bottlenecked populations (Bosse *et al*, 2012; Kardos *et al*, 2018). All models in the severe bottleneck scenario also had considerable variability among simulation runs. This variability is presumably due to the unpredictable effect that genetic drift can have on a population following a bottleneck. In the case where little diversity is preserved through a bottleneck, followed by inbreeding, ROH number will increase genome-wide. In another scenario, if a wider range of genetic diversity is preserved, the ROH number remains lower.

We also show population history interacting with selection, with little evidence for the effect of strong selection in the severe bottleneck scenario compared to a strong effect on ROH in the no-bottleneck scenario. This is a nice demonstration of the relationship between effective population size and efficiency of selection, in which populations experiencing a bottleneck show weaker responses to selection (Falconer, 1983; Kardos *et al*, 2021). In a large non-bottlenecked population there is increased genetic variation and selection is more efficient (Kardos *et al*, 2021; Petit and Barbadilla, 2009), allowing the beneficial alleles to become fixed and in doing so generate ROH hotspots. In contrast, in a smaller bottlenecked population, such beneficial alleles may be lost during the bottleneck and instead ROH hotspots can be the result of drift and inbreeding.

### Recombination rate

To investigate the effect of recombination rate on ROH in our dataset we used two maps, physical and genetic, which affected the number of ROHs detected and their location (Tables 1 & 2; Figure 2). In the Rum population three ROH hotspots found using the physical map were no longer classed as hotspots when using the genetic map. In addition, we found that some hotspots reached further over the hotspot threshold in the genetic map than in the physical map (Figure 2). The differing results are due to the negative correlation between recombination rate and ROH density, which is known from previous studies (Bosse *et al*, 2012; Pemberton *et al*, 2012).

In contrast to expectation and past literature (Bosse *et al*, 2012; Kardos *et al*, 2017), the addition of varied recombination rate in our simulation did not yield results that differed significantly from a neutral model (Figure 4). However, other models simulating the effect of recombination rate used higher values for recombination, to reflect the organism studied (Kardos *et al*, 2017). In red deer, the maximum recombination rate recorded anywhere in the genome is 4cM/Mb and the average across 32 acrocentric autosomes is 1.038cM/Mb (Johnston *et al*, 2017). In our simulations, the maximum recombination rate was 1.75 cM/Mb based on the average acrocentric values. By using representative values for our study organism we may not have seen such a dramatic effect of recombination rate as other studies. However, as discussed above, accounting for variable recombination rate did affect the ROH hotspots. Perhaps for this population, recombination may be an aid to the formation of ROH hotspots, but not the primary source.

### Selection

A number of studies have identified ROH hotspots to be sites of positive selection (Grilz-Seger *et al*, 2019; Peripolli *et al*, 2018; Purfield *et al*, 2017; Sabeti *et al*, 2002; Shihabi *et al*, 2022). When accounting for the influence of recombination rate, the red deer population on Rum showed five genomic regions with a ROH in more than 15% of the population (Figure 2). It is possible these regions may be sites of positive selection. However, we are cautious to draw this conclusion. A relatively low number of individuals in the total population contained a ROH at a hotspot, in comparison to other ROH hotspot studies, which are as high as >80% in livestock and ~20% in humans (Pemberton *et al*, 2012; Purfield *et al*, 2012; Purfield *et al*, 2017) – although direct comparison between studies is difficult due to the minimum ROH length employed. Instead, the regions we identify here may be sites generated by genetic drift. Therefore, we further assessed the likelihood that these ROH hotspots are caused by selection or drift.

From our simulations, we conclude that the majority of the ROH hotspots observed are unlikely to be caused by strong selection. Inclusion of strong selection in the Rum population scenario increased the ROH hotspot threshold over that in the empirical dataset (Figure 4A). We cannot, however, conclude the ROH hotspots are purely a result of genetic drift, as simulations suggest that ROH hotspots are equally likely to occur as a result of neutral processes as under weak selection (Figure 4A)

We tentatively suggest that the maximum peaks in ROH density in the Rum data (i.e. the two highest ROH hotspots on chromosomes 15 and 28, Figure 2) *could* be the result of selection. First, there is a sharp decrease in haplotype diversity at the onset of these two ROH hotspots (Figure 3). It can be assumed that positive selection leads to a sudden decrease in haplotype diversity (Shihabi *et al*, 2022), as certain haplotypes are being favoured. This assumption is also supported by our simulations. We saw multiple drops in haplotype diversity in the strong selection Rum simulation, some of which coincide with ROH hotspots, but we saw very few haplotype diversity drops in the neutral simulation. Second, these hotspots are also apparent when using only short ROH, between 2.5-5Mb (Supplementary Figure 4). Short ROH reflect more ancient inbreeding where IBD segments are broken down by recombination over time (Kirin *et al*, 2010; McQuillan *et al*, 2008). Hence, short ROH have been subject to selection pressures for a longer period of time and are more likely to show evidence for long-term selection (Zhang *et al*, 2015). Finally, the empirical value for the maximum ROH density (Peak ROH density, Figure 2) overlap values generated from the strong selection simulation (Figure 4B. Middle panel). However this empirical value was also within the range generated from the neutral or weak selection simulations.

The strength of selection certainly plays an important role in the formation of a ROH hotspot. Increasing the selection coefficient (*s*) in our simulations resulted in an increase in ROH density. This was particularly evident in the non-bottlenecked population (as high N_e_ allows for a stronger response to selection). By having high *s*, these new alleles are likely to go to fixation and generate a ROH hotspot. However, in artificially selected organisms *s* is much higher than the value we chose for our simulations (mean s=0.005). Therefore, in these populations with very high selection coefficients, selection is likely to be the driver of the majority of ROH hotspots (Kim *et al*, 2013). In our bottlenecked simulation, it is likely that genetic drift outweighed the strength of selection, as we see little response to the increase in *s*. Thus in populations with low Ne, ROH hotspots may instead be sites of genetic drift rather than selection. A very recent estimate suggests the Rum population has a N_e_ below 200 (Gauzere *et al*, 2022), and likely has lower values for *s* than artificially selected organisms. Therefore, the low N_e_ together with the findings discussed here suggest that most ROH hotspots in this population likely emerged due to drift, but we cannot exclude the possibility that some hotspots are the result of strong selection.

### Conclusion

Our simulations have offered a valuable way to assess how different factors may be affecting the distribution of ROH in our study population, and highlight the danger of assuming all ROH hotspots are sites of selection. Overall, simulations show that population history has the most substantial effect on ROH number and density. Empirical data supports the effect of population history, with particular influence on the number of ROH. We also show in our simulations that when N_e_ is high, and/or selection is particularly strong, selection is the main driver of ROH hotspots. However, in populations with weak selection, genetic drift has the main impact on the formation of ROH hotspots. Simulations show no effect of recombination rate; however, empirical data suggests otherwise. This suggests that recombination rate may aid the formation of ROH hotspots in our study population but is not the sole cause. In our study population conclude that the most likely driver of ROH formation is genetic drift, aided by genome-wide recombination. However, two ROH hotspots are also consistent with a strong selection scenario, but we can neither fully exclude drift as a driver.

## Acknowledgements

We thank NatureScot for permission to work on the Isle of Rum. We thank the many field workers that have helped to collect samples, particularly Alison and Sean Morris. The long-term project and this research were funded by the UK Natural Environment Research Council, and most of the SNP genotyping was supported by a European Research Council Advanced Grant to J. M. Pemberton. A. Hewett is supported by an E4 NERC DTP studentship (NE/S007407/1). M. Stoffel is funded by The Leverhulme Trust (RPG-2019-072). S.E. Johnston is supported by a Royal Society University Research Fellowship (UF150448). The Argyll sample genotypes were contributed by Eryn McFarlane. Genotyping was conducted at the Wellcome Trust Clinical Research Facility Genetics Core. We are grateful to the Darwin Tree of Life project for the physical map positions used. We thank Ino Curik and two anonymous reviewers for their helpful comments on this manuscript.

## Author contribution statement

J.M.P coordinated sampling and genotyping. A.M.H conducted analyses and drafted the manuscript. L.P conducted the haplotype phasing. M.A.S, L.P., S.E.J and J.M.P helped with analyses and interpretations of results.

All authors contributed to revisions.

## Conflict of interest

The authors declare no competing interests in relation to the work described.

## Data availability

Data will be archived upon acceptance.

## Notes

### Competing Interest Statement

The authors have declared no competing interest.

