## Supplementary figures and tables for "Selection, recombination and population history effects on runs of homozygosity (ROH) in wild red deer (*Cervus elaphus*)"

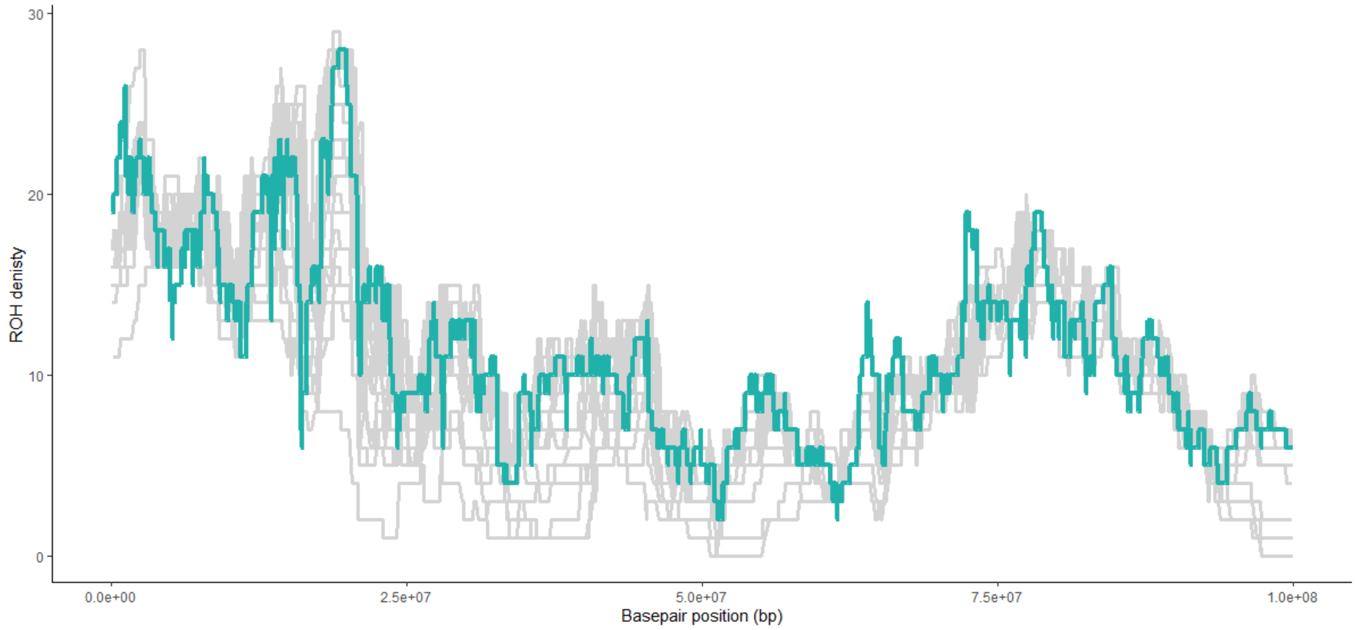


**Supplementary Figure 1** – True ROH density (coloured teal) in one iteration of a simulated chromosome using all possible existing SNPs (93748 SNPs in a 100Mb region) and searching for ROH using *PLINK* default search parameters. This is compared to 30 iterations in which SNPs were randomly thinned to the average SNP density used in our study (1 SNP every 61.2Kb or 1634 SNPs across a 100Mb region, shown in grey lines), and using *PLINK* parameters described in the main paper Methods section. The thinned iterations track the ROH density in the 'true ROH', indicating that our ROH density estimators is representative of the true ROH density in the population. The simulations conditions were the Rum population history scenario under neutral selection, see methods for more details.

**S2A)**

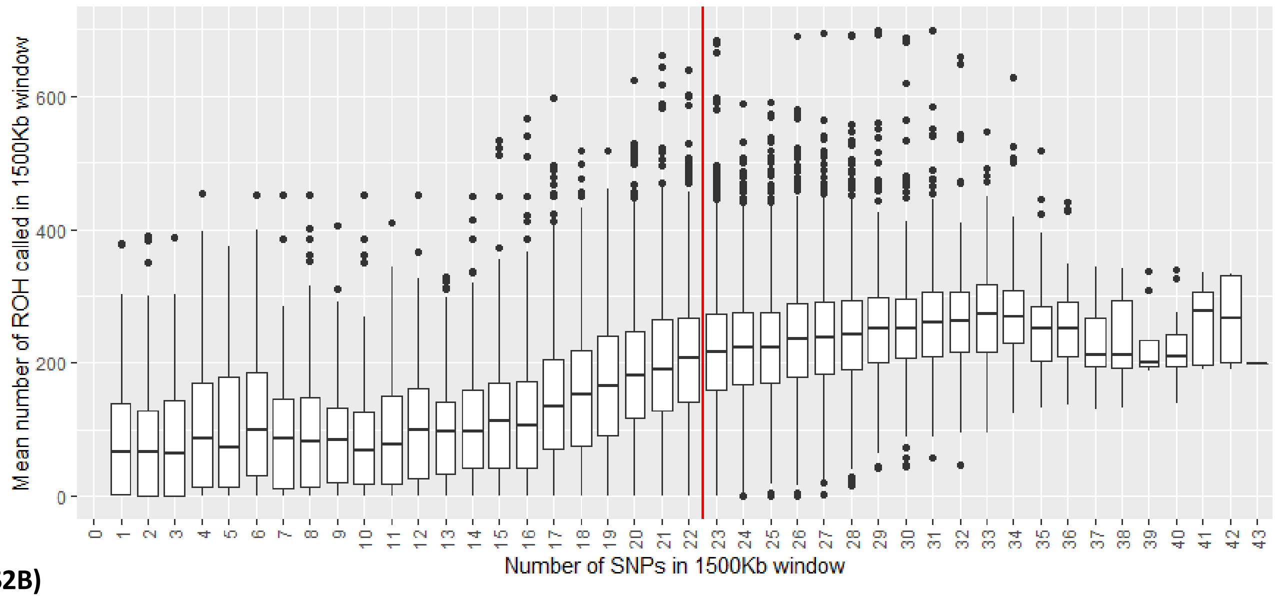

Mean number of ROH called in 1500Kb window
0
200
400
600
0 1 2 3 4 5 6 7 8 9 10 11 12 13 14 15 16 17 18 19 20 21 22 23 24 25 26 27 28 29 30 31 32 33 34 35 36 37 38 39 40 41 42 43
Number of SNPs in 1500Kb window


**S2B)**

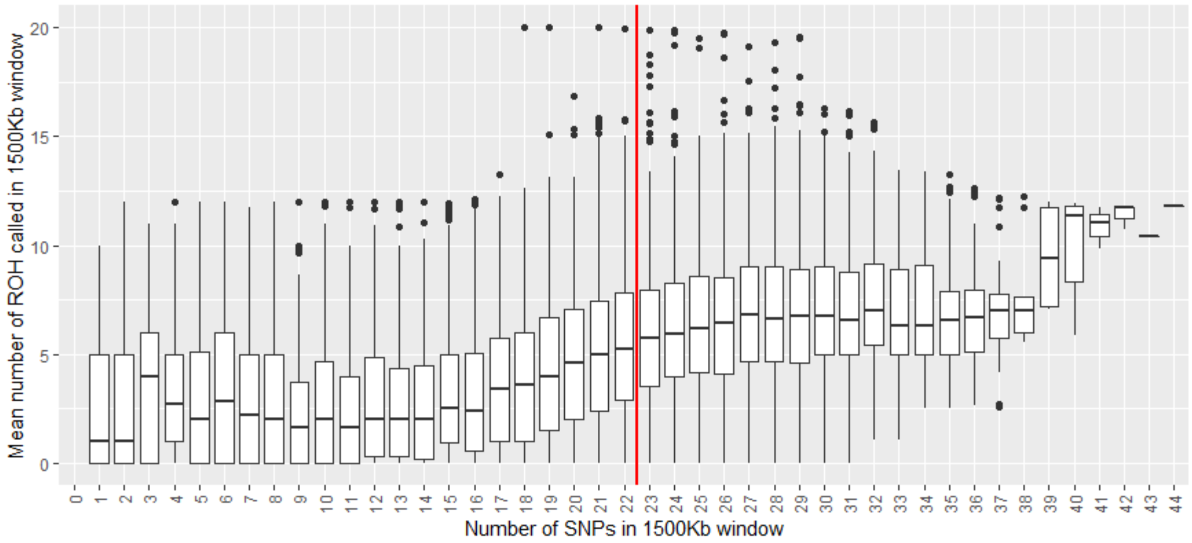

Mean number of ROH called in 1500Kb window
0
5
10
15
20
0 1 2 3 4 5 6 7 8 9 10 11 12 13 14 15 16 17 18 19 20 21 22 23 24 25 26 27 28 29 30 31 32 33 34 35 36 37 38 39 40 41 42 43 44
Number of SNPs in 1500Kb window

**S2C)**

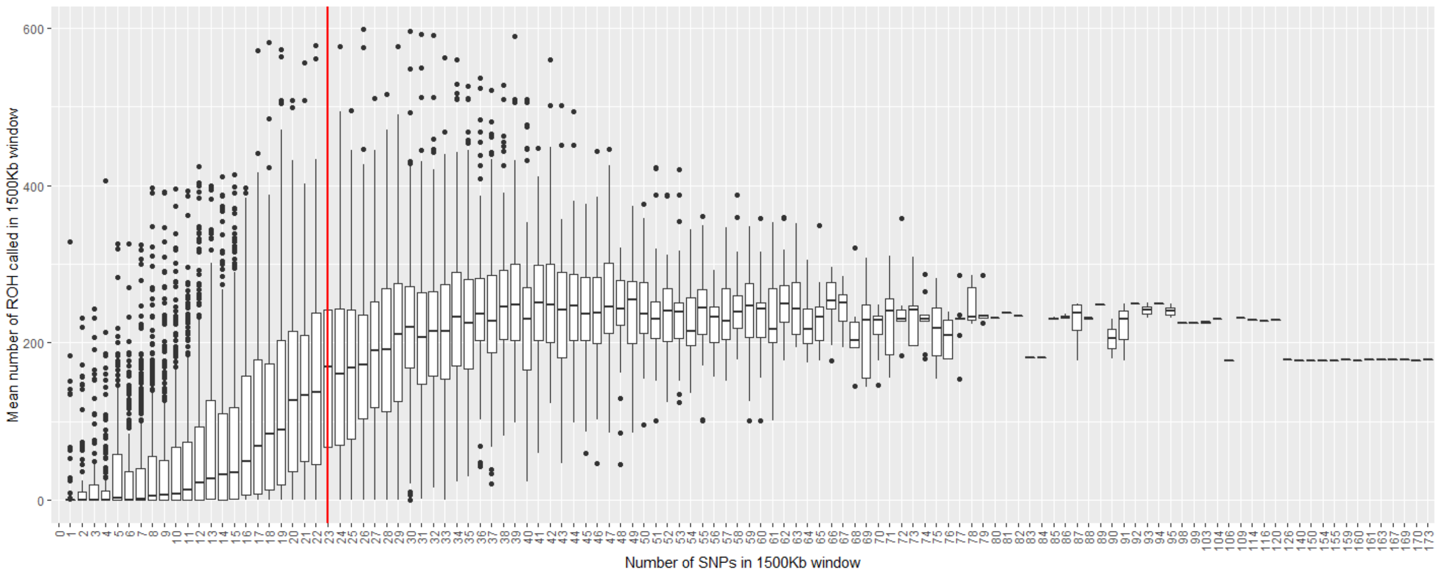


**S2D)**

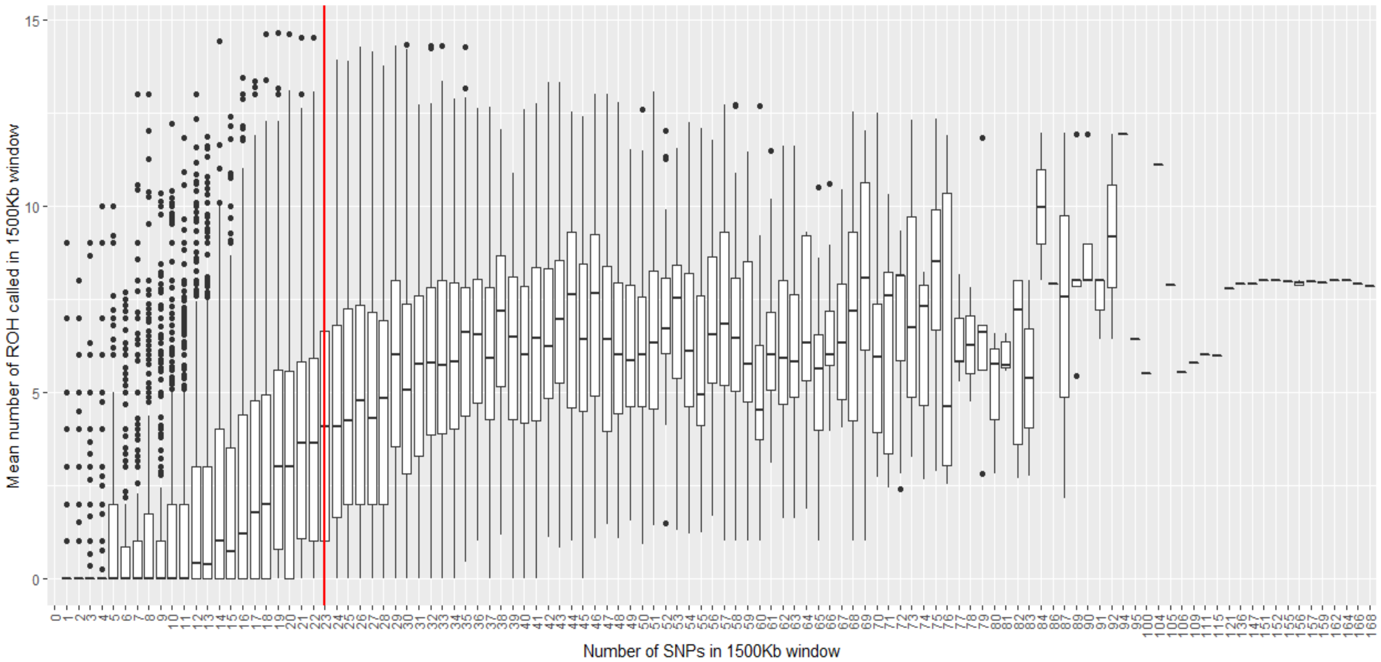


**Supplementary Figure 2** – Boxplots showing the relationship between number of SNPs within a 1500Kb window and the mean number of ROH called in same window in **S2A)** the Rum population using the physical (bp) map **S2B)** the Argyll population using the physical map **S2C)** the Rum population using the genetic (cM) map and **S2D)** the Argyll population using the genetic map. Solid red line indicates the cut off threshold for quality control steps post-ROH search. A minimum of 23 SNPs per 1500Kb window was chosen as an acceptable threshold value, all SNPs in windows below this value (to the left of the solid red line) were discarded.

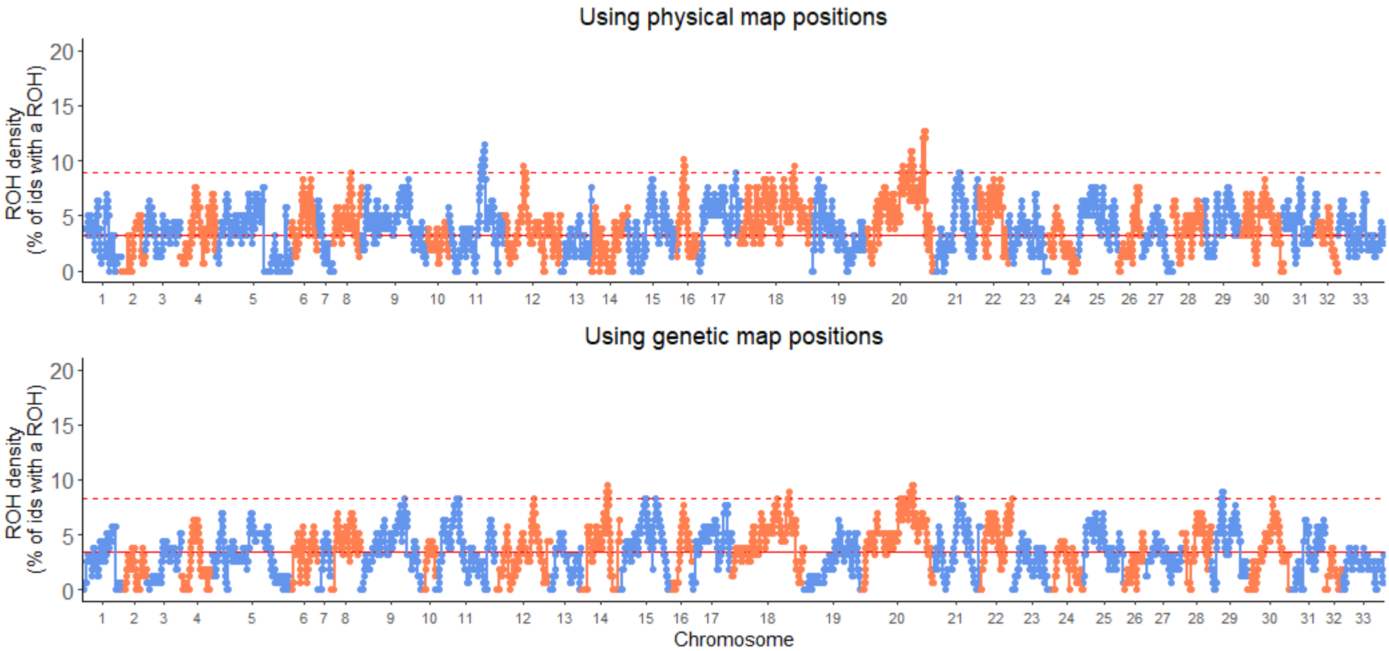


**Supplementary Figure 3** – Percentage of Argyll individuals with a ROH at each SNP (ROH density) across all chromosomes. The top panel shows the distribution of ROH density using the physical map positions (bp) and the bottom panel shows the same when using genetic map positions (cM). The upper dashed lines indicate the ROH hotspot threshold (99$^{th}$ percentile ROH density) and the solid red line indicates the mean ROH density.

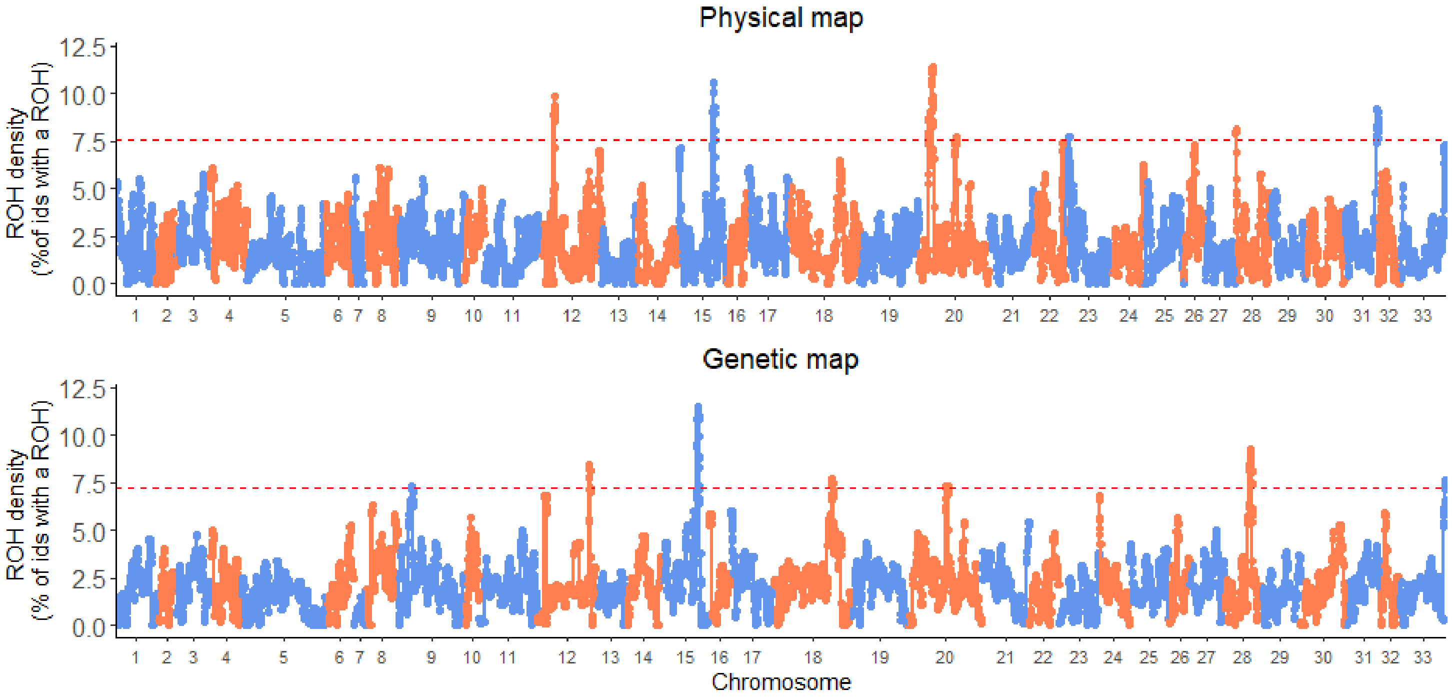


**Supplementary Figure 4** – Percentage of Rum individuals with a short ROH (2.5-5 Mb) at each SNP (ROH density) across all autosomal chromosomes. The top panel shows the distribution of ROH density using the physical map positions (bp) and the bottom panel shows the same when using genetic map positions (cM). The upper dashed lines indicate the ROH hotspot threshold (99$^{th}$ percentile ROH density) and the solid red line indicates the mean ROH density.

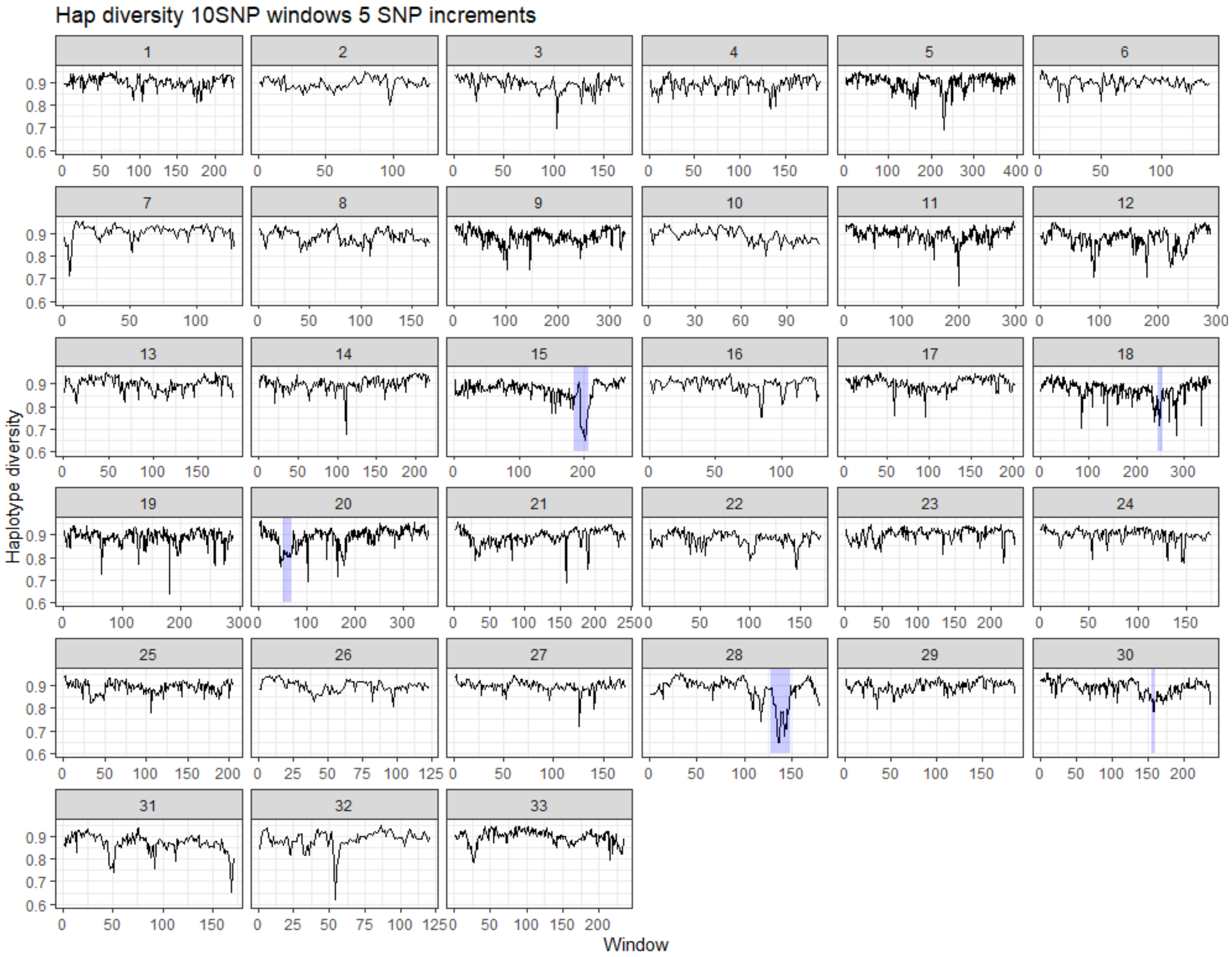


**Supplementary Figure 5** – Haplotype diversity for **10-SNP windows in 5 SNP sliding increments** across 33 autosomes in the Rum population of red deer. Low haplotype diversity shows certain haplotypes are more common, high haplotype diversity shows haplotypes occur in equal frequencies. ROH hotspot regions are shaded in purple.

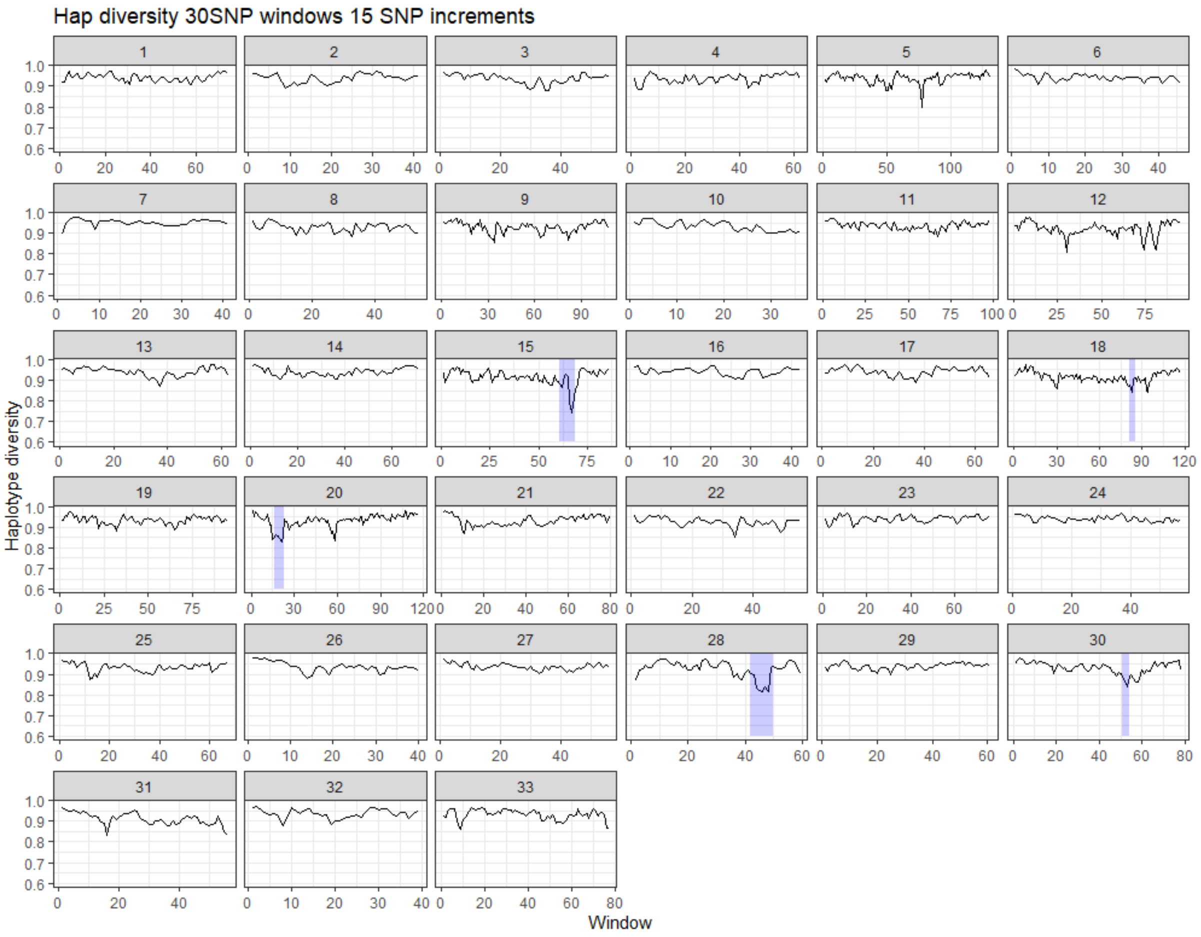


**Supplementary Figure 6** – Haplotype diversity for **30-SNP windows in 15 SNP sliding increments** across 33 autosomes in the Rum population of red deer. Low haplotype diversity shows certain haplotypes are more common, high haplotype diversity shows haplotypes occur in equal frequencies. ROH hotspot regions are shaded in purple.

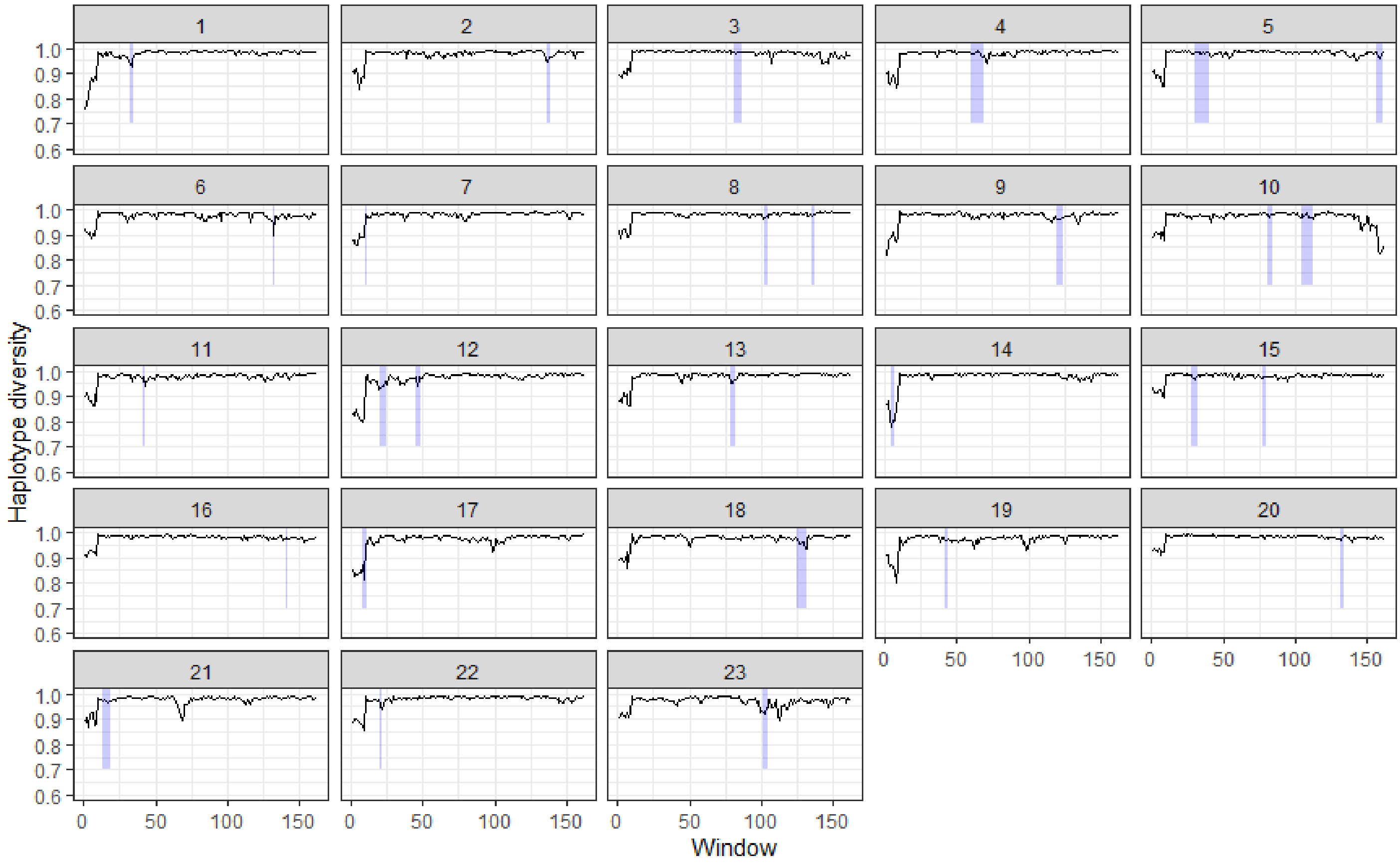


**Supplementary Figure 7** – Haplotype diversity in **neutral** simulations for the Rum scenario for 20-SNP windows in 10 SNP sliding increments across 23 simulated 100Mb chromosomes, details in main text. Raw SLiM outputs were thinned to 1630 SNPs per 100Mb region (average ROH density in empirical data), see supplementary Figure 1 for more details. Low haplotype diversity shows certain haplotypes are more common, high haplotype diversity shows haplotypes occur in equal frequencies. ROH hotspot regions are shaded in purple. NB: 2 simulation iterations failed to save raw outputs due to an error.

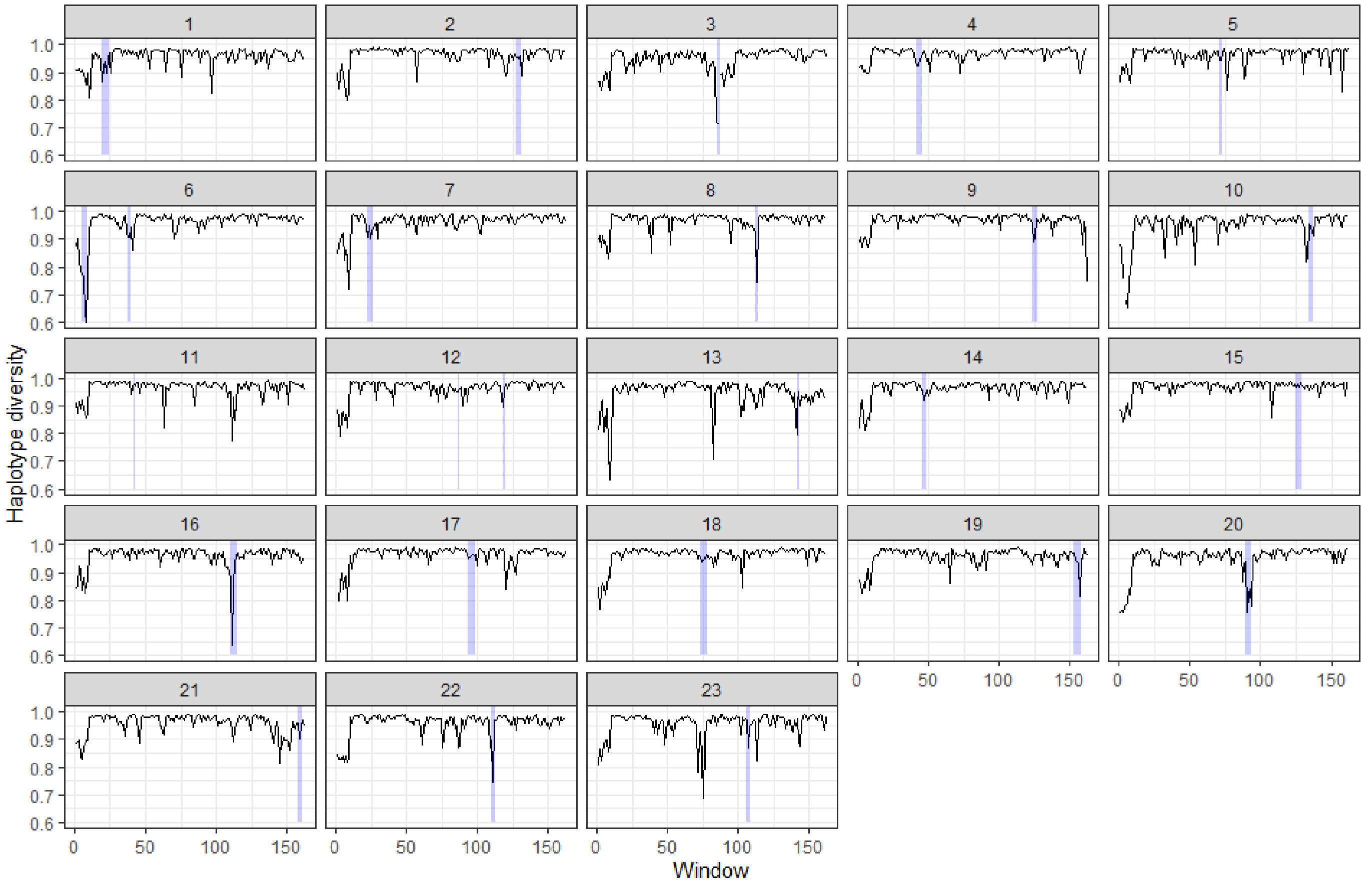


**Supplementary Figure 8** – Haplotype diversity in simulations with strong selection for the Rum scenario for 20-SNP windows in 10 SNP sliding increments across 23 simulated 100Mb chromosomes, details in main text. Raw SLiM outputs were thinned to 1630 SNPs per 100Mb region (average ROH density in empirical data), see supplementary Figure 1 for more details. Low haplotype diversity shows certain haplotypes are more common, high haplotype diversity shows haplotypes occur in equal frequencies. ROH hotspot regions are shaded in purple.

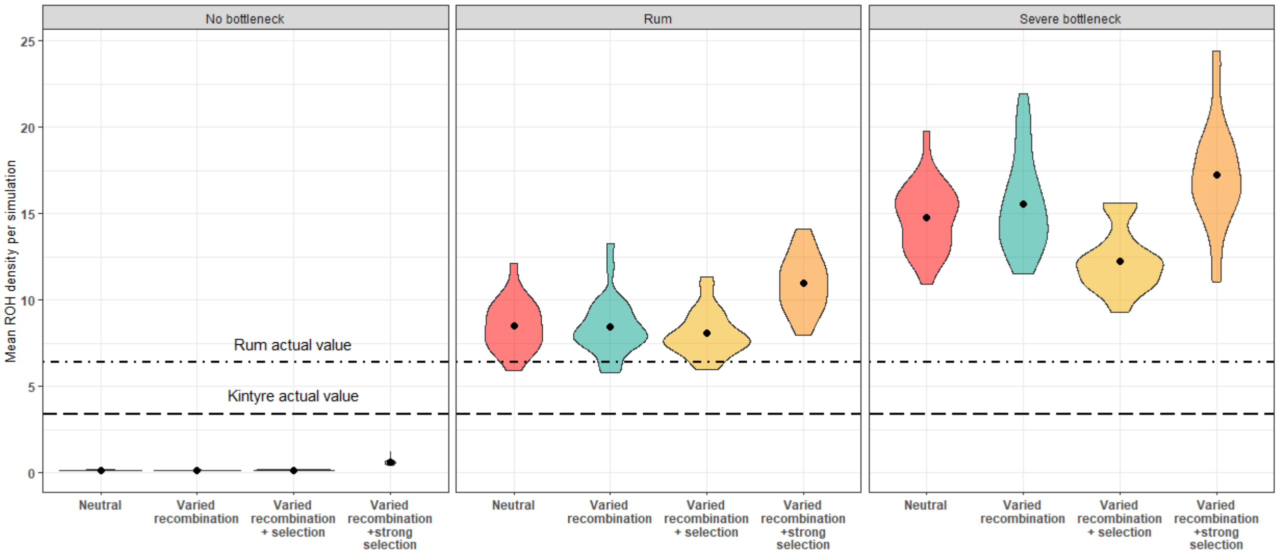


**Supplementary Figure 9 -** Violin plots showing mean ROH density for every simulation iteration. Black dots indicate the mean value for 23 iterations of a simulation, the width of the violin indicates the number of simulations at that value. Each box contains a different simulated population history, from left to right: no historical population bottleneck, Rum population history, severe historical population bottleneck. Each population history was modelled under four conditions: a neutral model with a constant recombination rate and no selection, a model including varied recombination rate, a model including varied recombination and selection and a model including varied recombination rate and strong selection (for details see main text). Top and bottom dashed lines show the same value for the Rum and Argyll empirical dataset respectively.

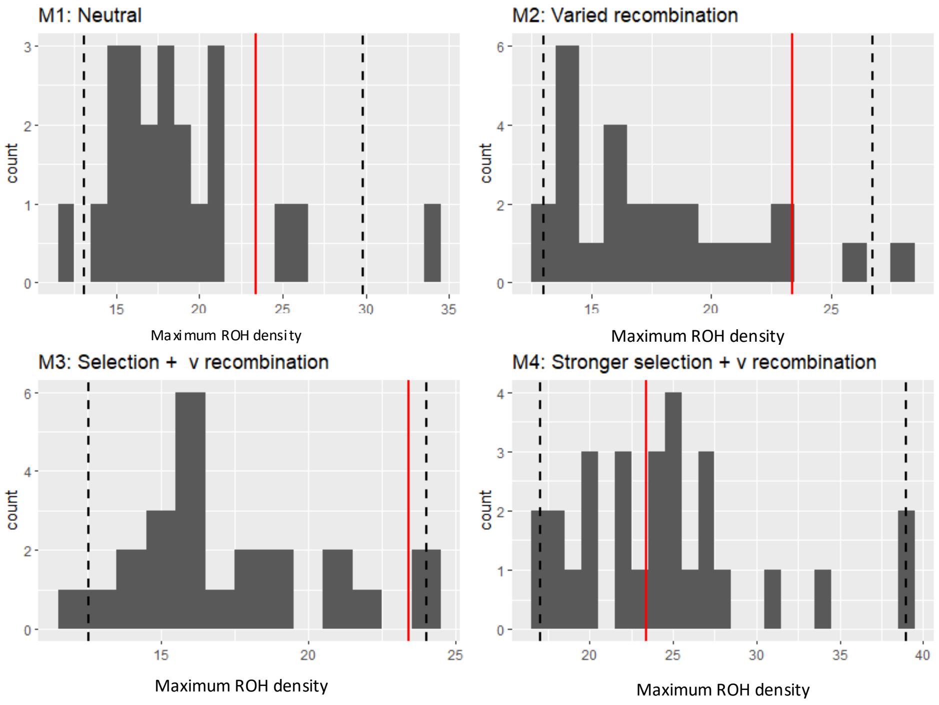


**Supplementary Figure 10** – Histogram of maximum ROH density values from 23 iterations of a simulated 100Mb chromosome for four different models under the Rum population history scenario. Model 1 (M1) is a neutral model, M2 is a model with varied recombination rate across the chromosome, M3 has an addition of weak selection and M4 has instead an addition of strong selection. Dashed lines show the 95% confidence intervals for maximum ROH density values across all 25 iterations. Solid red line shows the maximum ROH density value for the empirical Rum dataset.

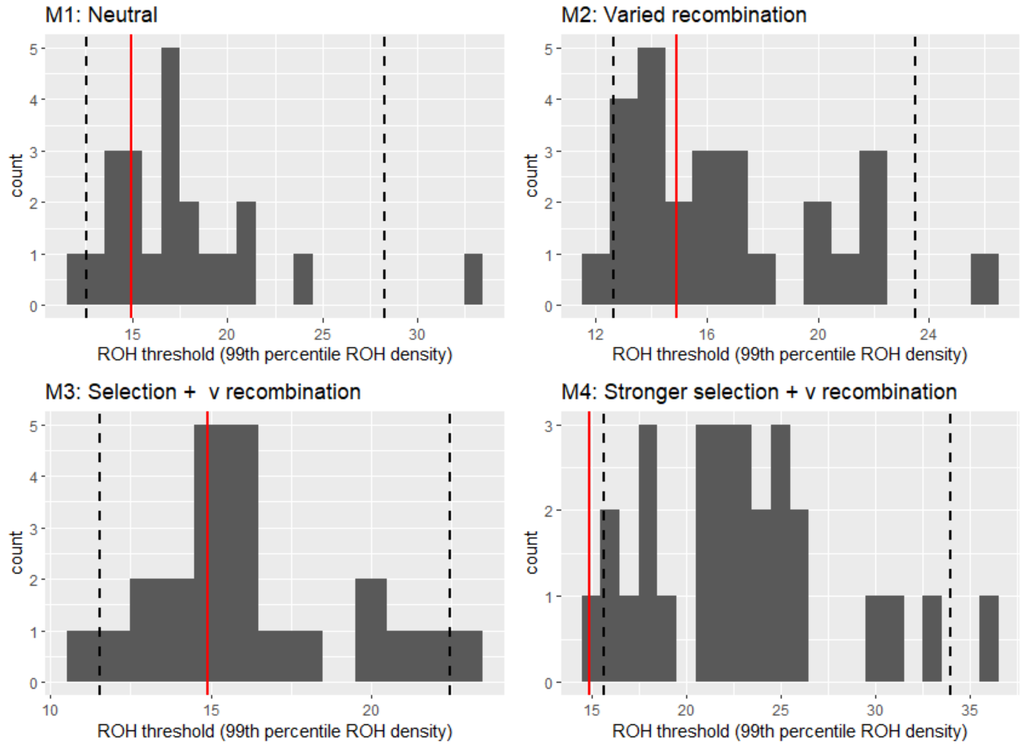


**Supplementary Figure 11 –** Histogram of ROH hotspot threshold values from 23 iterations of a simulated 100Mb chromosome for four different models. Model 1 (M1) is a neutral model, M2 is a model with varied recombination rate across the chromosome, M3 has an addition of weak selection and M4 has instead an addition of strong selection. Dashed lines show the 95% confidence intervals for ROH hotspot threshold values across all 25 iterations. Solid red line shows the ROH hotspot threshold value for the empirical Rum dataset

### Supplementary Methods

##### *ROH size classes and inbreeding coefficients*

Identified ROH were classified into three length categories based on the relationship between ROH length and generations since a common ancestor, $l$ = 100 2$g$ $cM$ (Thompson 2013). The categories were as follows, with the corresponding mean number of generations to the common ancestor in brackets:

Short ROH: 2.5mbp (~20 g) – 5mbp (10 g),
Medium ROH: 5Mbp (10g)– 16Mbp (~3 g),
Long ROH > 16Mbp (≤3 g)

The % of runs, mean number per id and proportion of the genome covered in each category were enumerated.

### Supplementary tables

**Supplementary table 1** – Pearson correlations between the number of SNPs and the number of called ROH in a 1500kn window before and after quality control (QC). Remaining SNPs show the number of SNPs used for analysis of ROH hotspots after QC.

| Data set | Pearson's correlation before density QC | Pearson's correlation after density QC | Remaining SNPs after density QC |
|---|---|---|---|
| **Rum bp positions** | 0.44 | 0.16 | 28,875 |
| **Rum cM positions** | 0.56 | 0.20 | 25,798 |
| **Kintyre bp positions** | 0.38 | 0.14 | 28,055 |
| **Kintyre cM positions** | 0.53 | 0.21 | 25,194 |

**Supplementary Table 2** – Table showing the percentage of total ROH, mean number per individual and the average percentage of the genome in a ROH for each ROH category using genetic and physical map positions for the Rum population . ROH categories include: short (2.5mbp – 5mbp), medium (5Mbp – 16Mbp), long (> 16Mbp) and all ROH.

| Population | Map used | ROH category | % of all ROH | Mean number of ROH per id | Mean $F_{ROH}$ for each length class |
|---|---|---|---|---|---|
| Rum | Genetic (cM) | ROH short | 55.8 | 8.4 | 1.21 |
| | | ROH medium | 40.4 | 6.08 | 1.94 |
| | | ROH long | 3.8 | 0.56 | 0.54 |
| | | ROH total | N/A | 15.04 | 3.70 |
| | Physical (bp) | ROH short | 52.6 | 9.5 | 1.37 |
| | | ROH medium | 41.9 | 7.6 | 2.50 |
| | | ROH long | 5.5 | 1.00 | 1.00 |
| | | ROH total | N/A | 18.2 | 4.88 |
| Kintyre | Genetic (cM) | ROH short | 58.4 | 4.6 | 0.66 |
| | | ROH medium | 37.2 | 2.9 | 0.97 |
| | | ROH long | 4.5 | 0.4 | 0.36 |
| | | ROH total | N/A | 7.8 | 1.94 |
| | Physical (bp) | ROH short | 57.5 | 5.3 | 0.75 |
| | | ROH medium | 36.7 | 3.4 | 1.11 |
| | | ROH long | 5.8 | 0.5 | 0.57 |
| | | ROH total | N/a | 9.2 | 0.02 |

**Supplementary Table 3** - Comparison of ROH hotspots using two maps to search for ROH in the **Kintyre** deer dataset. The table shows the mean ROH density, with standard error and the ROH hotspot threshold. Any SNP with a value of ROH density above this threshold value is classed as a ROH hotspot SNP. The table also shows the number of SNPs classed as a ROH hotspot SNP and the chromosomes containing ROH hotspots. The notation (c) indicate independent hotspots on the same chromosome as found in the Rum dataset.

| **Map Position used** | **Mean *ROH density* (%)** | **ROH Hotspot threshold (99$^{th}$ percentile *ROH density*)** | **# of SNPs over ROH hotspot threshold** | **Chromosomes containing ROH hotspots** |
|---|---|---|---|---|
| Physical | 3.7 ± 0.01 | 8.9% | 404 | 8,11,12,16,17,18(c),20(c),21 |
| Genetic | 3.4 ±0.02 | 8.2% | 92 | 14,18(c),20(c),29 |

**Supplementary table 4** – Details of genetic location of ROH hotspots in two study populations using physical and genetic map positions to search for ROH. CHR indicates the chromosomes number of the ROH hotspot with notations detailing unique positions. Start and end positions of ROH hotspots are given in basepairs (bp). Start and end SNPs detail the first and last SNP respectively included in the ROH hotspot. Equal position in cM indicates the equivalent position of that start and end SNPs identified when using the genetic map to search for ROH, hence is only applicable for the genetic map.

| Population | Map used | CHR | Start position in bp | Equal positions in cM | Start SNP | End position in bp | Equal position in cM | End SNP |
|---|---|---|---|---|---|---|---|---|
| Rum | Physical | 5 | 46651775 | n/a | cela1_red_17_24410527 | 48032482 | n/a | cela1_red_17_22899306 |
| | | 15 | 69172359 | n/a | cela1_red_26_26804781 | 72277338 | n/a | cela1_red_26_30124007 |
| | | 18(a) | 37934322 | n/a | cela1_red_4_28722688 | 37934322 | n/a | cela1_red_4_28722688 |
| | | 18(b) | 85716613 | n/a | cela1_red_4_79357066 | 91784187 | n/a | cela1_red_4_85897274 |
| | | 20(a) | 48218637 | n/a | cela1_red_3_24187404 | 50696346 | n/a | cela1_red_3_26770667 |
| | | 20(b) | 81465370 | n/a | cela1_red_3_59627920 | 84697925 | n/a | cela1_red_3_63056189 |
| | | 28 | 46600920 | n/a | cela1_red_9_45485811 | 51403457 | n/a | cela1_red_9_50865789 |
| | | 30 | 54779069 | n/a | cela1_red_12_56171583 | 54779069 | n/a | cela1_red_12_56171583 |
| | Genetic | 15 | 66041717 | 63401000 | cela1_red_26_22794294 | n/a | 69822000 | cela1_red_26_30528628 |
| | | 18 | 89025258 | 68056000 | cela1_red_4_82863021 | 91784187 | 70006000 | cela1_red_4_85897274 |
| | | 20 | 43931654 | 30584000 | cela1_red_3_21530324 | 51074650 | 35097000 | cela1_red_3_27166017 |
| | | 28 | 46551039 | 49956000 | cela1_red_9_45442363 | 51664874 | 54040000 | cela1_red_9_51142731 |
| | | 30 | 54684785 | 57101000 | cela1_red_12_56063030 | 55341642 | 57584000 | cela1_red_12_56796140 |
| Kintyre | Physical | 8 | 33603732 | n/a | cela1_red_2_101452924 | 33912322 | n/a | cela1_red_2_101136015 |
| | | 11 | 64560919 | n/a | cela1_red_11_39796647 | 70199360 | n/a | cela1_red_11_33805906 |
| | | 12 | 35792781 | n/a | cela1_red_10_66406560 | 38683715 | n/a | cela1_red_10_63272849 |
| | | 16 | 16993389 | n/a | cela1_red_8_96805514 | 19127728 | n/a | cela1_red_8_94277412 |
| | | 17 | 55442786 | n/a | cela1_red_6_58005078 | 55697563 | n/a | cela1_red_6_58287621 |
| | | 18 | 92990164 | n/a | cela1_red_4_87210156 | 93761012 | n/a | cela1_red_4_88012599 |
| | | 20 | 85199216 | n/a | cela1_red_3_63610534 | 90100608 | n/a | cela1_red_3_68857519 |
| | | 20 | 96254044 | n/a | cela1_red_3_75438618 | 100973704 | n/a | cela1_red_3_80489601 |
| | | 20 | 113909537 | n/a | cela1_red_3_94225098 | 117503504 | n/a | cela1_red_3_97977175 |
| | | 21 | 55788800 | n/a | cela1_red_14_56080868 | 56549886 | n/a | cela1_red_14_56934289 |
| | Genetic | 14 | 55075434 | 52715000 | cela1_red_16_54570687 | 57680072 | 54427000 | cela1_red_16_57487234 |
| | | 18 | 93761012 | 72122000 | cela1_red_4_88012599 | 93761012 | 72122000 | cela1_red_4_88012599 |
| | | 20 | 96391617 | 66136000 | cela1_red_3_75578861 | 100973704 | 70770000 | cela1_red_3_80489601 |
| | | 29 | 25547950 | 30089000 | cela1_red_8_25976505 | 26010655 | 30866000 | cela1_red_8_26504836 |

**Supplementary table 5** – Table showing summary information for SLiM simulations for 3 different population histories and 4 different models tested within each history.

| Population | Model | Mean number of ROH per id | Average KB in a ROH per id |
|---|---|---|---|
| No bottleneck | **Model 1: neutral** | 0.02692 | 129.9 |
| | **Model 2: recombination** | 0.027136 | 129.8 |
| | **Model 3: selection & recombination** | 0.032717 | 152.4 |
| | **Model 4: strong selection & recombination** | 0.17241 | 627.7 |
| Rum | **Model 1: neutral** | 1.64 | 8576.8 |
| | **Model 2: recombination** | 1.61 | 8485.8 |
| | **Model 3: selection & recombination** | 1.47 | 8178.9 |
| | **Model 4: strong selection & recombination** | 2.17 | 11124 |
| Bottleneck | **Model 1: neutral** | 3.32 | 15009.1 |
| | **Model 2: recombination** | 3.36 | 15772.6 |
| | **Model 3: selection & recombination** | 2.56 | 12406 |
| | **Model 4: strong selection & recombination** | 3.76 | 17447.1 |
